# ACES: Analysis of Conservation with Expansive Species

**DOI:** 10.1101/2021.06.16.448733

**Authors:** Evin M. Padhi, Elvisa Mehinovic, Eleanor I. Sams, Jeffrey K. Ng, Tychele N. Turner

## Abstract

**Motivation:** An abundance of new reference genomes are becoming available through large-scale sequencing efforts. While the reference FASTA for each genome is available, there is currently no automated mechanism to query a specific sequence across all new reference genomes.

**Results:** We developed ACES (Analysis of Conservation with Expansive Species) as a computational workflow to query specific sequences of interest (e.g., enhancers, promoters, exons) against reference genomes with an available reference FASTA. This automated workflow generates BLAST hits against each of the reference genomes, a multiple sequence alignment file, a graphical fragment assembly file, and a phylogenetic tree file. These data files can then be used by the researcher in several ways to provide key insights into conservation of the query sequence.

**Availability:** ACES is available at https://github.com/TNTurnerLab/ACES

**Supplementary information:** Supplementary Figure 1 is available on bioRxiv.

## 1 Introduction

Recently, long-read sequencing and other advanced genomic technologies have enabled cost-effective reference genomes in several species (Rhie et al., 2021). Exemplar projects, such as the Vertebrate Genome Project (Rhie et al., 2021), have an ambitious goal of generating high-quality reference genomes for 66,000 different vertebrate species. This powerful dataset will provide key insights in numerous areas. An area of interest in the study of human disease is to assess the conservation of specific elements in the genome (e.g., enhancers, promoters, exons), especially those that lack a well-defined genetic code.

In performing characterization of various genomic elements, we sought to use these new reference genome resources to gain deeper insights about our elements of interest. However, there was no quick way to query these sequences of interest against the new reference genomes resulting in a bottleneck in our work.

To alleviate this problem, we developed an automated workflow to query sequences across all reference genomes and ultimately generate a multi-sequence alignment file, graphical alignment file, and phylogenetic tree files.

One interesting example, which doubled as a positive control for the workflow, that we focus on in the present study is the ZRS enhancer that targets the *SHH* gene and is involved in preaxial polydactyly (Lettice et al., 2003). This element is highly conserved in vertebrates but has lost its function in snakes due to a 17 nucleotide deletion in a specific transcription factor binding site that has been functionally validated to modulate limb development (Kvon et al., 2016).

## 2 Application

Analysis of Conservation with Expansive Species (ACES) is a computational workflow that performs evolutionary analysis of a given query sequence, in FASTA format, by comparing it to genome data from any reference genomes of interest to the user. Visualization of the workflow is shown in Supplementary Figure 1.

ACES starts by taking a sequence of interest and performs a BLASTn (version 2.5.0+) (Altschul et al., 1990) search on each genome present in the database. Using a search strategy designed to identify distantly related orthologs, it returns sequences meeting user defined e-value, length and percent identity thresholds while simultaneously recording the hits in a log file. Next, ACES performs a multiple sequence alignment using MUSCLE (v3.8.1551) (Edgar, 2004) to generate both a multiple sequence alignment file and PHYLIP output files. Then, the multiple sequence alignment file is run through a python program (our modified version of https://github.com/fawaz-dabbaghieh/msa_to_gfa version 1.0.0) to convert the multiple sequence alignment file to a graphical fragment assembly (GFA) file. A GFA file is a tab-delimited text file that describes sequences and their overlaps. With the help of an outside GFA viewer (e.g., BANDAGE (Wick et al., 2015)), the user can import the generated GFA file to view the sequence graph and visualize conservation. Finally, ACES utilizes RAxML (version 8.2.12) (Stamatakis, 2014) to generate a phylogenetic tree file and estimate the rates of evolution at the locus of interest. By default, ACES uses the GTRGAMMA model of evolution and performs bootstrap for Maximum Likelihood (ML) trees. Once finished, the tree file with the best likelihood is generated along with four supporting RAxML files that can be used in downstream evolutionary analysis.

## 3 Results

As an example, we describe herein our testing of ACES on a sequence located within the ZRS enhancer region (https://www.ncbi.nlm.nih.gov/gene?Db=gene&Cmd=DetailsSearch&Term=105804841). Specifically, we tested a 766 nucleotide subset of this sequence that had previously been examined and shown to contain a 17 nucleotide deletion in snakes (Kvon et al., 2016). When this deletion is introduced in mice they have a reduced limb growth phenotype. We sought to determine whether our fully automated approach could recapitulate the results in the Kvon paper. We aggregated the same 18 reference genomes, as found in the Kvon paper, and utilized our default settings of a BLAST e-value threshold of 0.00001 and a query length fraction of at least 0.3. After all the files were generated by our program, including a multiple sequence alignment file, best maximum likelihood tree file, and a GFA file, we proceeded to characterize the output files using existing downstream programs.

First, we assessed the multiple sequence alignment in MEGA (Kumar et al., 2016) and observed the expected 17 nucleotide deletion in snakes (Figure 1A). We then examined the best tree and found that the snakes all cluster together (Figure 1B) and also represented this in a BANDAGE plot using the GFA file to show that most of the element is conserved aside from a few small regions (Figure 1C). Overall, this demonstrated our workflow was working correctly.

**Figure 1:**
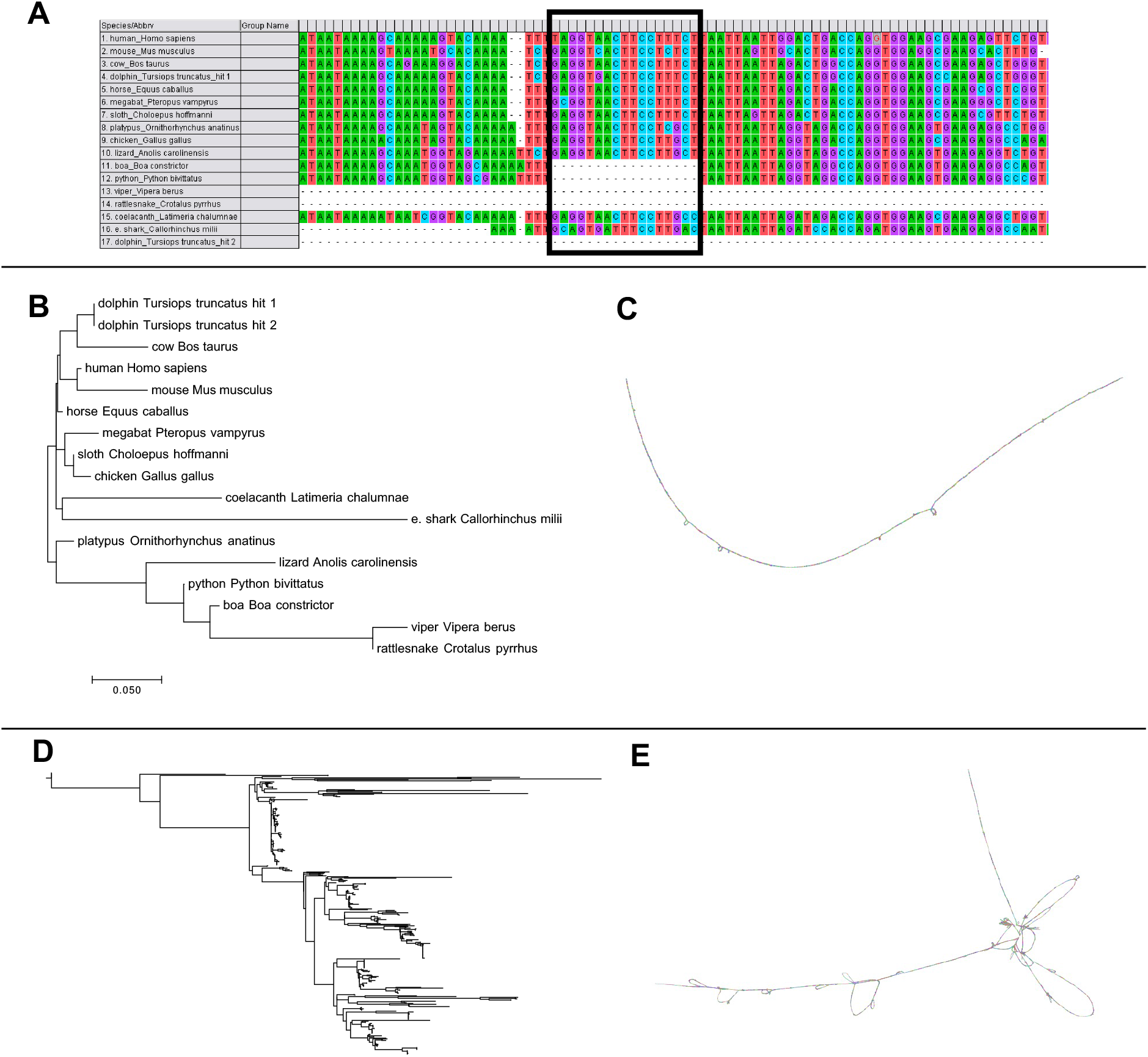
Testing ACES using a fragment of the ZRS enhancer. Figure 1A, 1B, and 1C all assess the same 18 reference genomes as in Kvon et al. 2016. Figures 1D and 1E assess the 522 reference genomes currently available in the Vertebrate Genome Project and Ensembl. A) The multiple sequence alignment output from ACES (visualized in MEGA) identifies the 17 nucleotide snake-specific deletion in this element; B) The phylogenetic tree from ACES shows a snake-specific section in the tree; C) The GFA file from ACES (visualized in BANDAGE) shows an overall high degree of conservation in the element; D) A more extensive phylogenetic tree generated from running ACES across all 522 currently available reference genomes; E) Detailed assessment of the ZRS sequence by examining the GFA across all 522 currently available reference genomes.

To further demonstrate the utility of our pipeline, we performed a comprehensive analysis of the same sequence using all the currently available Vertebrate Genome Project and Ensembl genomes (n = 522). This ran within two hours and we again examined the output data to see the full phylogenetic tree (visualized using icytree.org in Figure 1D). The BANDAGE plot of this analysis revealed a more complex diagram of the conservation of this element (Figure 1E). On average, our branches in the phylogenetic tree file had a bootstrap value of 62.5 with running 100 bootstraps.

## 4 Discussion

Reference genome data is being generated rapidly and we are on the path to new biological discoveries with this diverse data. We identified a current gap caused by the new data, which is the ability to utilize all of this data to quickly look at specific regions of interest. To address this gap, we developed ACES as a fast workflow to query sequences of interest and derive a multiple sequence alignment file, best maximum likelihood tree file, and GFA file for each sequence of interest. ACES is flexible and allows for testing of any number of reference genomes, which will be a boon as many new genomes continue to become available in the coming years. This workflow will be of interest to individuals assessing small sequences (e.g., enhancers, promoters, exons) across many genomes and useful in identifying new regions for downstream analyses and functional investigation.

## Acknowledgements

Thank you to members of the Turner Laboratory at Washington University in St. Louis for helpful discussions on this work.

## Data Availability

Reference genomes are publicly available at the Ensembl website at http://ftp.ensembl.org/pub/ and the specific versions used in this paper are also available on the Turner Lab Public Globus Endpoint at https://app.globus.org/file-manager?origin_id=97668938-bcc8-11eb-9d92-5f1f6f07872f&origin_path=%2F.

