## Supplementary figures and images for "ACES: Analysis of Conservation with Expansive Species"

### Supplementary_Figure_1

#### Supplementary Figure 1: ACES computational workflow.

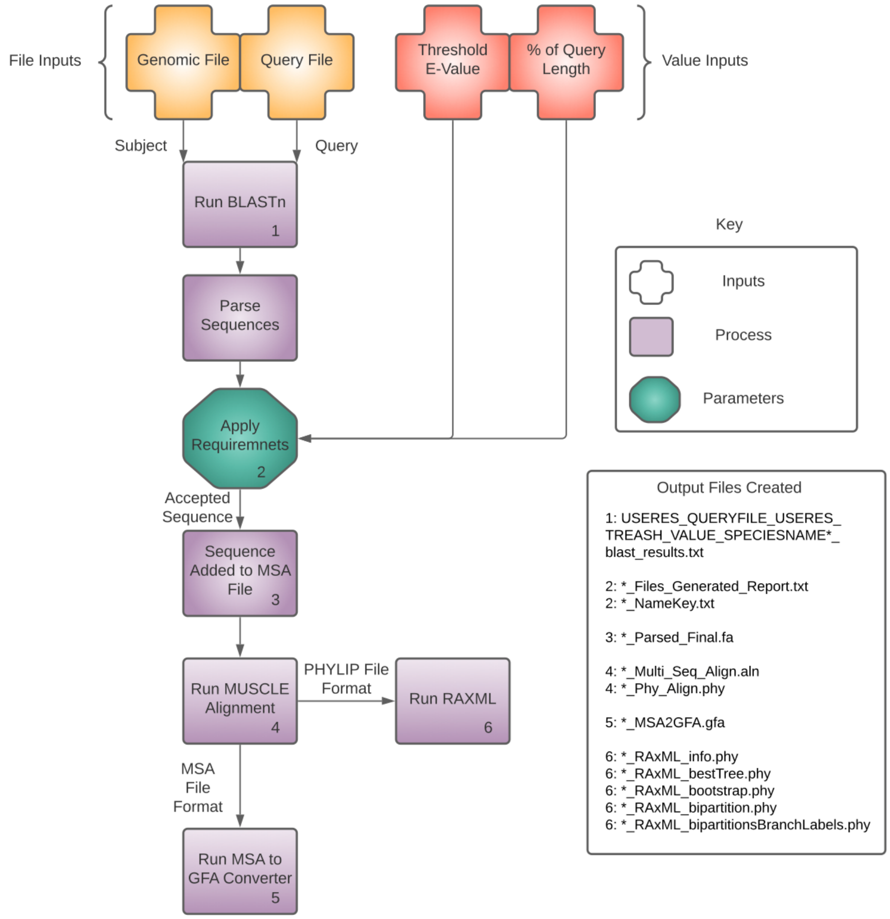
